# A Novel High-Throughput Single B-Cell Cloning Platform for Isolation and Characterization of High-Affinity and potent SARS-CoV-2 Neutralizing Antibodies

**DOI:** 10.1101/2022.03.20.485024

**Authors:** Paritosh Parashar, B Prabhakar, Sonali Swain, Nisha Adhikari, Punit Aryan, Anupama Singh, Mohit Kwatra

## Abstract

Monoclonal antibodies (mAbs) that are specific to SARS-CoV-2 can be useful in diagnosing, preventing, and treating the coronavirus (COVID-19) illness. Strategies for the high-throughput and rapid isolation of these potent neutralizing antibodies are critical toward the development of therapeutically targeting COVID-19 as well as other infectious diseases. In the present study, a single B-cell cloning method was used to screen SARS-CoV-2 receptor-binding domain (RBD) specific, high affinity, and neutralizing mAbs from patients’ blood samples. An RBD-specific antibody, SAR03, was discovered that showed high binding (ELISA and SPR) and neutralizing activity (competitive ELISA and pseudovirus-based reporter assay) against Sars-CoV-2. Mechanistic studies on human cells revealed that SAR03 competes with the ACE-2 receptor for binding with the RBD domain (S1 subunit) present in the spike protein of Sars-CoV-2. This study highlights the potential of the single B cell cloning method for the rapid and efficient screening of high-affinity and effective neutralizing antibodies for Sars-CoV-2 and other emerging infectious diseases.

**Highlights:**

1. Single B-cell cloning is a high-throughput and efficient method of generating high affinity neutralizing antibodies
2. Single B-cell cloning method was used to screen SARS-CoV-2 receptor-binding domain (RBD) specific, high affinity, and neutralizing monoclonal antibodies from patient’s blood samples.
3. An RBD-specific antibody, SAR03, was discovered that showed high binding and neutralizing activity against SARS-CoV-2.

## 1. Introduction

The severe acute respiratory syndrome coronavirus 2 (Sars-CoV-2) causes the coronavirus disease (COVID-19) that has crossed the species barrier and led to a global pandemic leading to high morbidity and mortality in humans along with significantly impacted on health, environmental, and socioeconomic status [1, 2]. At least 400 million cases of Sars-CoV-2 infections and 6 million deaths have been reported worldwide since the first case of COVID-19 was registered [3]. After the detection of the first Sars-CoV-2 variant in the Wuhan in the late year 2019, various mutants have emerged with higher infectivity rates compared to the original strain [4]. To effectively target the burden of COVID-19, the discovery and development of novel therapeutics, diagnostics, and vaccines are required. Encouragingly, several vaccine candidates against Sars-CoV-2 have emerged as timely prophylactic treatment with great success globally. However, the rapidly evolving variants of the new Sars-CoV-2 pose a threat to the COVID-19 vaccine effectiveness and hence, immediate development of alternate therapeutic interventions is needed. Although vaccine’s effectiveness in blocking infectious diseases and antibody therapy is an alternative treatment strategy in the prevention of newly emerging mutant strains of Sars-CoV-2.

Passive administration of high affinity neutralizing antibodies against SARS-CoV-2 may constitute an essential role in targeting COVID-19 and complement vaccine-based prophylactic intervention. Hybridomas and phage display techniques are among the most commonly used platforms for generating antibodies for research and therapeutic purposes [5, 6]. While the hybridoma method is limited with a longer time (6-8 months) for generating library, low efficiency, and requirement of humanization step, phage display is associated with high cost, generation of biased repertoires, and loss of natural pairing information [7, 8]. Rapid and efficient screening methods of identifying neutralizing mAbs against infectious diseases are in great demand.

In recent years, the emergence of Single B cell technologies has greatly accelerated the timelines for identification and further development of the natural repertoire of neutralizing antibodies against infectious diseases HIV, Ebola, and influenza [9–11]. This process involves the isolating PBMCs from recovered patient’s blood, high-throughput single-cell sorting of desired antigen-specific antibody expressing B-cells, recovery of the paired variable region of antibody’s heavy and light chain (V_H_/V_L_) genes through RT-PCR including two-step PCR steps, expression, and evaluation of antibody candidates. The application of single B-cell cloning has also allowed the identification of neutralizing mAbs against Sars-CoV-2, [12–16]. Several neutralizing mAbs have received emergency use approval for COVID-19 diagnosis, treatment, and even prophylactic use in patients exhibiting mild-to-moderate symptoms that reduces disease progression and subsequent hospitalization [17–19]. Sars-CoV-2 acquires entry within host cells and primarily interacts through the receptor-binding domain (RBD) (within the S1 subunit of spike protein) with the angiotensin-converting enzyme 2 (ACE-2) which is a cellular receptor [20, 21]. With the critical nature of the RBD interaction with ACE-2 for viral entry, antibodies with the potential ability to bind the RBD and interfere with ACE-2 binding can have potent neutralizing activity. Therefore, RBD emerges as the primary antigen target to specifically sort antibody expression B-cells against Sars-CoV-2.

This study aimed to evaluate the utility of single B cell sorting for the isolation of potent neutralizing antibodies against SARS-CoV-2. Here, we describe an optimized and rapid system for high-throughput screening of neutralizing mAbs through separating RBD antigenspecific IgG1^+^ memory B cells derived from the blood of patients recovered from Sars-CoV-2 infection. Through our optimized single B-cell cloning platform, we identified a neutralizing mAb, SAR03, with high affinity and neutralizing ability against Sars-CoV-2.

## 2. Materials and Methods

### 2.1 Determination of neutralizing antibodies in patient’s plasma

Blood from the 8 selected COVID-19 hospitalized but recovered donors were collected in EDTA tubes and plasma was collected for the presence of antibodies while the cells were further processed for B-cell sorting.

For binding ELISA, 1μg/mL recombinant RBD protein (in-house expressed) was coated to Nunc MaxiSorp ELISA plates (44-2404-21; Thermo) in carbonate buffer (pH-9.5) was kept overnight at 4°C. Thereafter, plates were proceeded with washing in TBST (1X TBS +.05% Tween-20) and blocked in 3% BSA prepared in TBST for 1 hr at room temperature. Afterward, the washing was done with TBST, followed by plasma addition to the plates at 1/10 dilution for 1 hr at 37°C. Further, washing was performed with TBST, with the addition of HRP conjugated Goat Anti-Human IgG (H+L) (109-035-088; Jackson ImmunoResearch) at 1/3000 dilution for 30 minutes at room temperature. Plates were then washed with TBST and TMB substrate (T0440; Sigma) was added for color development. The stop solution was added to terminate the reaction and absorbance was taken at 450 nm using a microplate reader.

For the neutralization activity of antibodies, competitive ELISA was used. 1μg/mL recombinant ACE-2 protein (in-house expressed) was coated onto the Nunc MaxiSorp ELISA plates (44-2404-21; Thermo) using carbonate buffer (pH-9.5) was kept overnight at 4°C. Subsequently, coated plates were washed in TBST (1X TBS +.05% Tween-20) and blocked in 3% BSA prepared in TBST for 1 hr at room temperature. Plasma at 1/10 dilution was pre-incubated with recombinant RBD-biotin (50ng/well) for 1 hr at 37°C in 100μl volume. The plasma-RBD mix was then added to ACE-2 coated plates and incubated for an additional 1 hr at 37°C. After washing the plates with TBST, HRP conjugated streptavidin (N100; Thermo) was added at 1/10000 dilution for 30 min incubated at room temperature.

The next step of washing was done in TBST and TMB substrate (T0440; Sigma) was added for color development. The stop solution was added to stop the reaction followed by absorbance measurement at 450 nm using a microplate reader.

### 2.2 Sorting of RBD-specific memory B cells by FACS

Isolation of blood mononuclear cells was performed using density gradient centrifugation (ACCUSPIN System-Histopaque-1077; Sigma) according to the manufacturer’s instructions. PBMCs from patients (showing binding and neutralization in plasma samples) were pooled and memory B-cells were isolated using Memory B Cell Isolation Kit (130-093-546; Miltenyi Biotech), as per the manufacturer’s protocol, using LD Columns and MidiMACS™ Separator. Cells were successively stained for 30 minutes on ice with the cocktail of fluorescently conjugated antibodies in 200 μl staining buffer (1X PBS + 2% FBS): 2 μg/ml biotinylated-RBD-Streptavidin-PE and FITC-anti-human IgG antibody (Biolegend, clone: M1310G05) and DAPI. Following washing of cells, the FACS analysis was achieved via BD FACSAria III (BD Biosciences) FSC-A versus SSC-A gating was performed for identification of total cell population, and RBD-specified single memory B cells were gated by IgG^+^RBD^+^ followed by single-cell sorting into 96-well PCR plates (free of DNase and RNase, Bio-Rad) containing 10μl lysis buffer (1X reverse transcriptase reaction buffer containing RNAse inhibitor). The plates were then stored at −80 °C for further processing.

### 2.3 Amplification, cloning, and expression of antibodies

We used an optimized heavy and light chain amplification method (in-house platform, unpublished data) where PCR primers were designed from different regions (leader, variable and constant regions) of immunoglobulin (Ig) as interpreted by the IMGT reference directory (http://www.imgt.org/vquest/refseqh.html). To begin with the first step in RT-PCR, 5uL μl of the RT-A mix was put into each well of 96 well plates containing a single B cell (prepare a master-mix according to the number of samples). Further, the mixture was incubated at 65°C for 5 min and immediately transferred to ice and kept for 3 min. 5 μl RT-B mix was added into each well of the plate with reaction program: 45 °C for 45 min, 70 °C for 15 min **(Table 1)**. Two replica plates were then prepared for heavy and light chains (kappa) by transferring 2μlof RT-PCR mix into new 96-well plates. 18μlof 1^st^ PCR mix was added to each well. The PCR program was scheduled as 1st PCR run: 95°C for 3 min, 30 cycles of 95°C for 10 sec, 55°C for 5 sec and 72°C for 1 min. 2 μl of the obtained 1st PCR product was then introduced into each well of a new 96 well plate holding 18μl 2^nd^ PCR mix. The PCR program for 2^nd^ run: PCR: 95°C for 3 min, 35 cycles of 95°C for 10 sec, 55°C for 5 sec, and 72°C for 45 sec. The amplification of 2^nd^ PCR product was analyzed through DNA gel electrophoresis. Samples showing the correct size band for heavy and light chain amplification in the respective wells were subsequently cloned into pAb20-hCHIgG1 (Synbio; for heavy chain) and pAb20-hCK (Synbio; for light chains) through in-fusion cloning (Takara). The ligated product was transformed to DH5-α competent cells, and the plasmid was purified through plasmid miniprep columns.

**Table 1:**
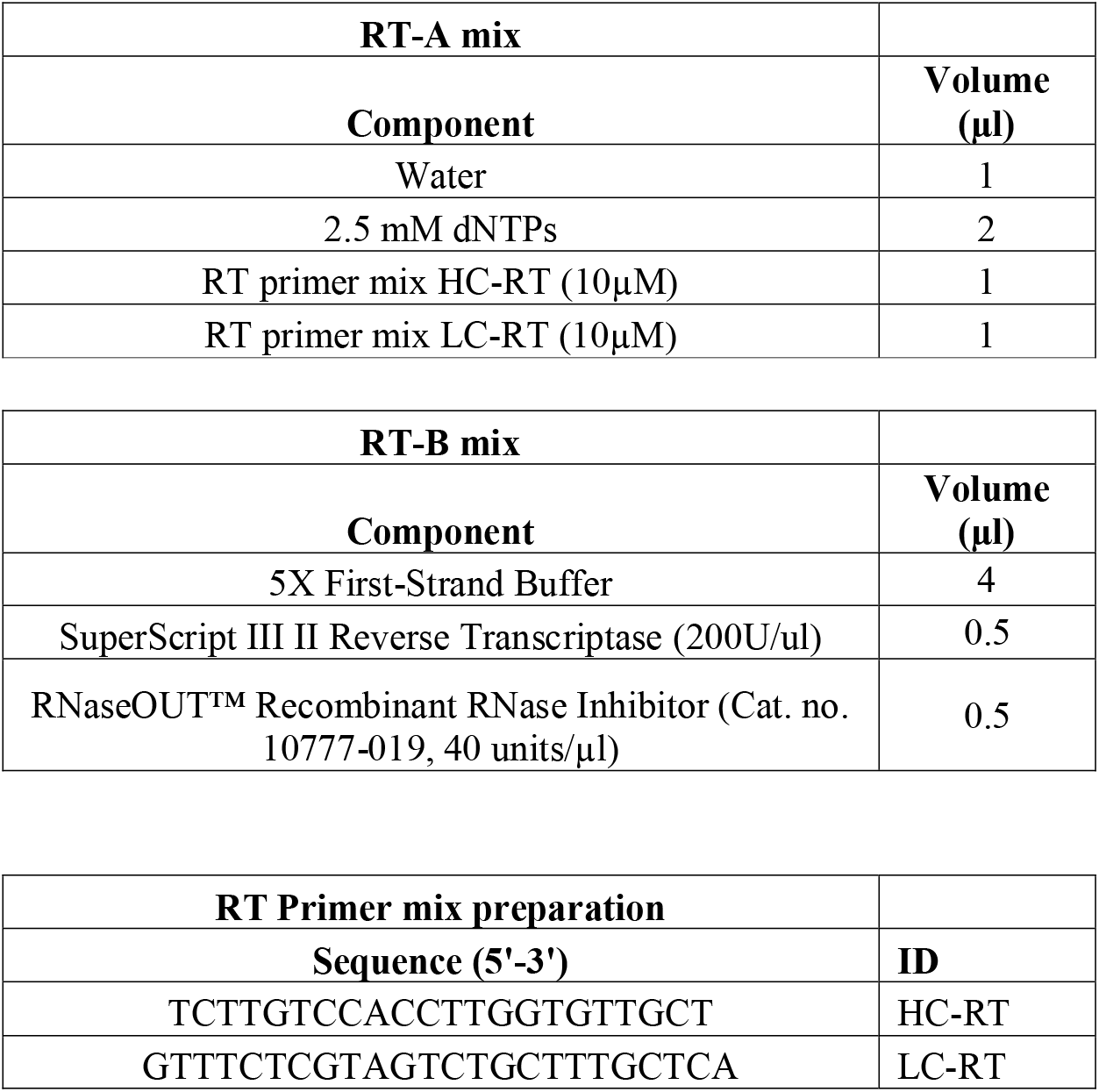
RT PCR details used in experiments: RT-A and RT-B mix compositions, Primer sequences details of HC-RT and LC-RT

Freestyle-293F cells (R79007; Thermo Fisher) were cultured in a 96-well culture plate with a density of 10000 cells/well. 24 hours later, pAb20-hCHIgG1 and pAb20-hCK plasmids discretely expressing heavy and light chain of antibodies were transiently co-transfected into Freestyle-293F cells (R79007, Thermo Fisher) employing purefection reagent (LV750A-1; System Bio) as per the instructions given by the manufacturer. Cells were further cultured for another 48 hours on condition of 5% CO2 at 37 °C. The supernatant was analyzed for secretion of antibodies through ELISA. Binding ELISA and neutralization activity of antibodies were performed as above. Cell culture supernatant was used at 1/10 dilution.

### 2.4 Stable cell line generation and recombinant antibody production

Freestyle-293F cells (R79007; Thermo Fisher) cells were grown in a 6-well culture plate at a density with 2×10^6^ cells/well/2ml media. 24 hours later, pAb20-hCHIgG1 and pAb20-hCK plasmids separately expressing heavy and light chain of antibodies were transiently cotransfected into Freestyle-293F cells (R79007; Thermo Fisher) using purefection reagent (LV750A-1; System Bio) followed by manufacturer protocol. 48-hours post-transfection, cells were selected in presence of hygromycin (200μg/ml) and blasticidin (10μg/ml) for the selection of stable clones expressing heavy and light chains. The selected individual colonies were observed for 2 weeks proceeded with trypsinization and limited dilution method implied for generation of single-cell monoclones. Furthermore, monoclones showing the highest titer in ELISA were selected for further characterization.

The stable cells were then cultured in a shaker incubator run at 120 rpm with conditions of 8% CO_2_ at 37 °C. After one week, the supernatants consisting of secreted antibodies were collected and trapped by protein G Sepharose (GE Healthcare). The bound antibodies on the Sepharose were eluted and concentrated using Centricon Plus-70 (051555; Millipore). Successively, purified antibodies were utilized in the following binding and neutralization analysis.

### 2.5 Antibody binding affinity measurement by surface plasmon resonance (SPR)

The affinity of antibody-binding recombinant RBD was assessed using the Biacore T200 platform. The CM5 chip (GE Healthcare) was attached with our novel Sars-CoV-2 antibody to record 6000 response units. Gradient concentrations of recombinant RBD protein were diluted (2-fold dilution, ranging from 50 nM −0.78 nM) using HBS-EP+ Buffer (0.01 M HEPES; 0.15 M NaCl; 0.003 M EDTA; and 0.05% (v/v) Tween-20, pH 7.4) and was allowed to flow through the chip surface. Every cycle, the sensor was recovered using Gly-HCl (pH 1.7). The antibody and RBD affinity ability were estimated employing a 1:1 binding fit model in Biacore Evaluation software (GE Healthcare).

### 2.6 Pseudovirus neutralization assay

Pseudovirus was created as described in an earlier report [22]. HEK293T cells were seeded in a T-75 flask at a density of 4.5×10^6^ cells/flask/15ml media. The next day plasmid DNA was diluted in 500μl OptiMEM at a ratio of Transfer vector (CMV-GFP-T2A-Luciferase, BLIV101PA-1, System-bio) viral packaging (psPAX2): viral envelope (pMD2G) or Sars-CoV-2 envelope (pMT1-SARS-CoV-2-S, custom gene synthesis; Synbio) at 4:2:1 ratio (6:3:1.5ug, respectively). 40μl Purefection was diluted in 500μl OptiMEM media. DNA and purefection were mixed followed by an incubation step of 15 minutes at room temperature and then added to the plated cells. The supernatants were isolated and collected after 48 hours proceeded with filtration by 0.45 μm membrane filter and then centrifugated at 300xg for 10 min. The obtained supernatant was aliquoted and then immediately stored at −80°C until further analysis.

For pseudovirus-based neutralization assay, HEK293 cells stably expressing ACE-2 receptor (HEK293-ACE2) were plated into a tissue culture treated opaque white 96-well microplate in complete medium at a density of 5000 cells/well. Cells were incubated at 37°C with 5% CO2 overnight. The next day, serial dilution of antibody at 10X final concentration in the complete medium was prepared and pre-incubated 10μl of antibody with 80μl pseudovirus for 30 minutes at room temperature. Antibody and pseudovirus suspension was then added to the HEK293-ACE2 cells and 10μl of 50μg/ml (10X) of polybrene to the cells (5μg/ml final concentration). The plate was centrifuged at 800rpm for 60 minutes. 72 hours after transduction, the rate of transduction was observed through visualizing GFP expression. Luciferase reagent was prepared (ONE-Glo Luciferase Assay System; E6110, Promega) per recommended protocol. Recommended volume of Luciferase Assay reagent was added per well. The plate was then incubated at room temperature for the recommended time and the luminescence was measured using a luminometer. Lastly, the half-maximal inhibitory concentrations (IC50) were measured using the four-parameter logistic regression in GraphPad Prism 8.0.

### 2.7 Statistical analysis

All grouped data are expressed as the mean ± standard deviation (SD) of a demonstrative experiment executed at least in triplicate, and almost similar data were obtained in at least three independent experiments. The statistical analyses were directed using GraphPad Prism 8.0. One-way ANOVA followed by Dunnet multiple comparison tests of individual data values were used for statistical analysis. The P < 0.05 was considered statistically significant.

## 3. Results

### 3.1 Sorting of RBD specific memory B-cells

To identify and clone Sars-CoV-2 neutralizing antibodies, we first obtained the plasma and PBMC from the blood of 8 convalescent patients infected with COVID-19. These isolated plasma samples were first tested for binding to recombinant RBD protein **(Fig.1A)** and neutralizing activity through competition ELISA **(Fig.1B and C).** Using binding ELISA, we confirmed that the antibodies present in the plasma of all the patients displayed a high binding affinity to recombinant RBD. Upon experiment for competitive ELISA, we observed that plasma samples of three patients showed high neutralizing activity (Patient 2, 3, and 5) **(Fig. 1C)**. With these findings, we decided to utilize these three samples for sorting of RBD-specific memory B-cells.

**Fig 1.**
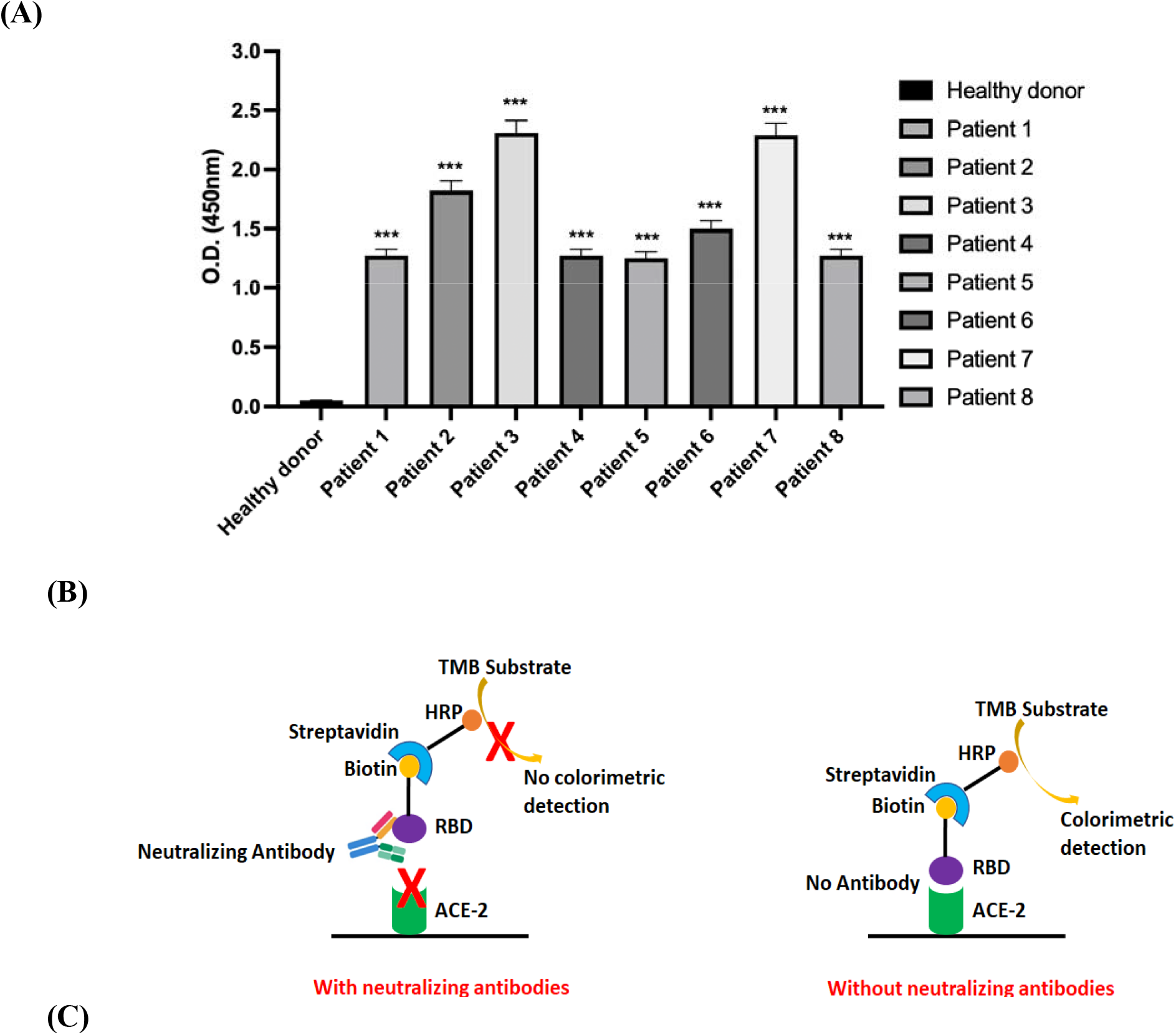
Screening of neutralizing antibodies against Sars-CoV-2 in patient’s plasma through ELISA. **(A)** Determination of antibody affinity through ELISA. The patient’s plasma was incubated in ELISA plates coated with recombinant RBD protein followed by detection with anti-Human-HRP antibody. **(B)** Schematic of competitive ELISA for checking neutralizing activity. C, Screening for neutralizing activity of antibodies present in patient’s plasma. Results are represented as Mean ± SD (n = 3). ***P<0.001 on comparison patient’s plasma with healthy donor plasma.

Since the SARS-CoV-2 virus principally infects human cells through molecular interaction of RBD domain to human cell surface ACE2 receptor, therefore the recombinant RBD was used as a bait to sort the specific memory B cells via flow cytometry. Pan-memory cells were first isolated using the memory-B cell isolation kit using magnetic separation. Isolated B-cells were then stained with anti-IgG-FITC and RBD-Biotin-Streptavidin-PE. RBD-specific B-cells were then gated for IgG^+^RBD^+^ cells **(Fig.2).** Finally, 83 RBD-specific memory B-cells were then sorted into a 96-well plate, one cell per well, for antibody gene isolation.

**Fig 2.**
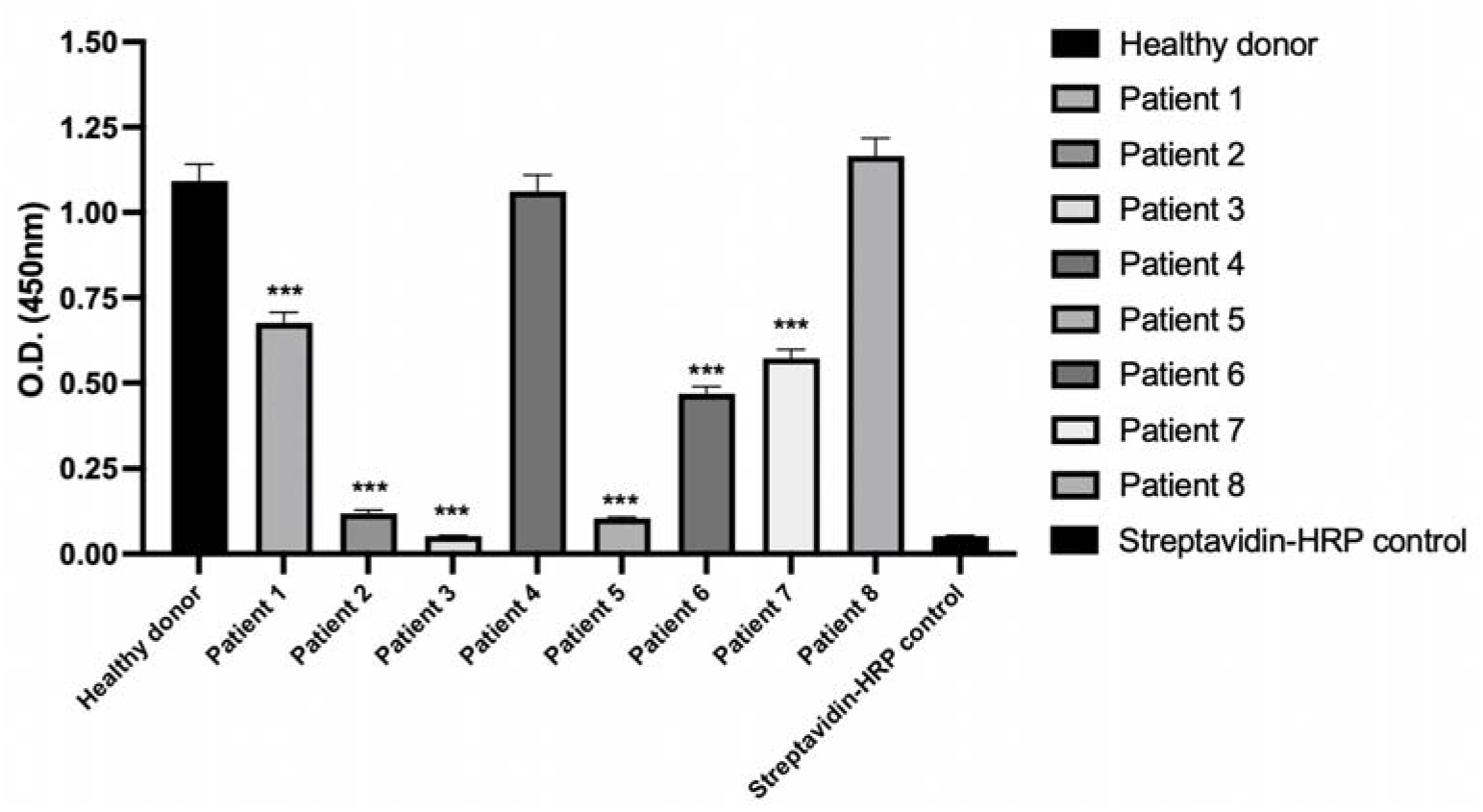
Isolation of RBD-specific memory B cells using flow cytometry. Live IgG+ (FITC conjugated) and RBD+ (PE-conjugated) cells were selected using single B-cell sorting and sorted as single cells in a 96-well plate.

### 3.2 Antibody heavy and light chain amplification and cloning

Immunoglobulin heavy and light (kappa) chains were acquired from the sorted single memory B-cells through RT-PCR and modified two-step nested PCR method **(Fig.3A)** [23, 24]. For RT-PCR, a mix of reverse primer from the constant region of the antibody sequence were used (HC-RT for heavy chain and LC-RT for kappa light chain, unpublished data). After RT-PCR, obtained cDNA was duplicated into two 96-well plates for amplification of heavy and light chain sequences separately. We utilized a mix of 20 nucleotides sequences located at the 5’ end of the native signal peptide in the antibody genes as the forward primers in the 1st PCR step. This forward primer was linked with a 15-bp adapter sequence that was overlapping with the 3’ end of the CMV promoter in the expression plasmid to be used in infusion cloning. HC-RT and LC-RT were again used as reverse primers for 1^st^ PCR used for amplification of heavy and light chain sequences, respectively. For the 2^nd^ PCR, the adapter sequence was used as forward primer and a mix of reverse primers for J-region of antibody gene along with 15-bp adapter sequences overlapping with 5’ sequence of the constant region already presents in pAB20 plasmids. Agarose gel was run to analyze the amplification of heavy and light chain sequences **(Fig.3B).** Out of 83 sorted single B-cell, 52 cells showed the amplification of both heavy and light chains with the correct size **(Fig.3B**, amplification of ten samples showed as the representative image for heavy and light chain amplification) and were further cloned into the expression plasmids. After 2^nd^ PCR, the linearized PCR product was cloned into linearized plasmid through in-fusion cloning utilizing 15-bp adapter sequence overlap. This high-throughput and improved method of amplification allowed us to complete the process of B-cell sorting to antibody sequence cloning in 7-10 days. Finally, 12 clones were obtained with heavy and light chains cloned in expression plasmids.

**Fig 3.**
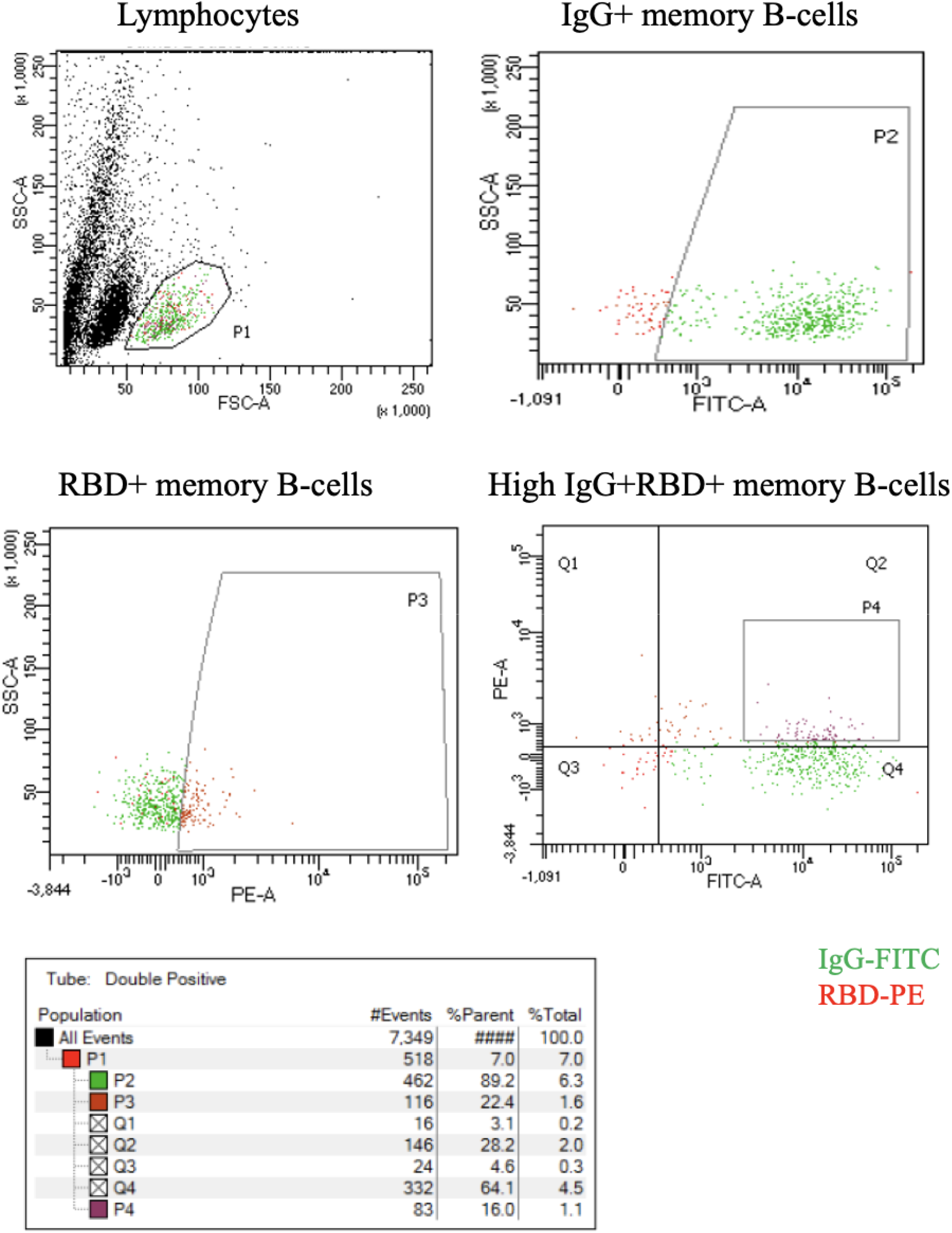
Method of cloning antibody heavy and light chain sequences. **(A)** Schematic of RT-PCR and PCR amplification of the variable region of heavy and light chain region of the antibody. **(B)** RT-PCR and 2-step PCR were carried out to amplify the variable region of heavy and light chain sequences along with their natural leader sequence. Amplified sequences were cloned into a pAb20-hCHIgG1 plasmid with hygromycin (for Gamma heavy chain) and pAb20-hCK plasmid with blasticidin (for kappa light chain).

### 3.3 High-throughput recombinant antibody production, binding analysis, and neutralization activity

Plasmid expressing heavy and light chains of the final 12 antibody clones were transfected into Freestyle-293F cells in a 96-well plate followed by supernatant collection after 48 hrs time interval. The supernatant was first analyzed for binding to recombinant RBD through ELISA. All 12 clones showed potent binding at 1/10 dilution of supernatant **(Fig.4A).** All the clones were then tested for neutralizing activity through competitive ELISA. Three clones (SAR03, SAR09, and SAR051) showed significantly good neutralizing activity (at 1/10 dilution of supernatant) **(Fig.4B)** and were selected for the generation of stable cell lines and large-scale protein production and purification.

**Fig 4.**
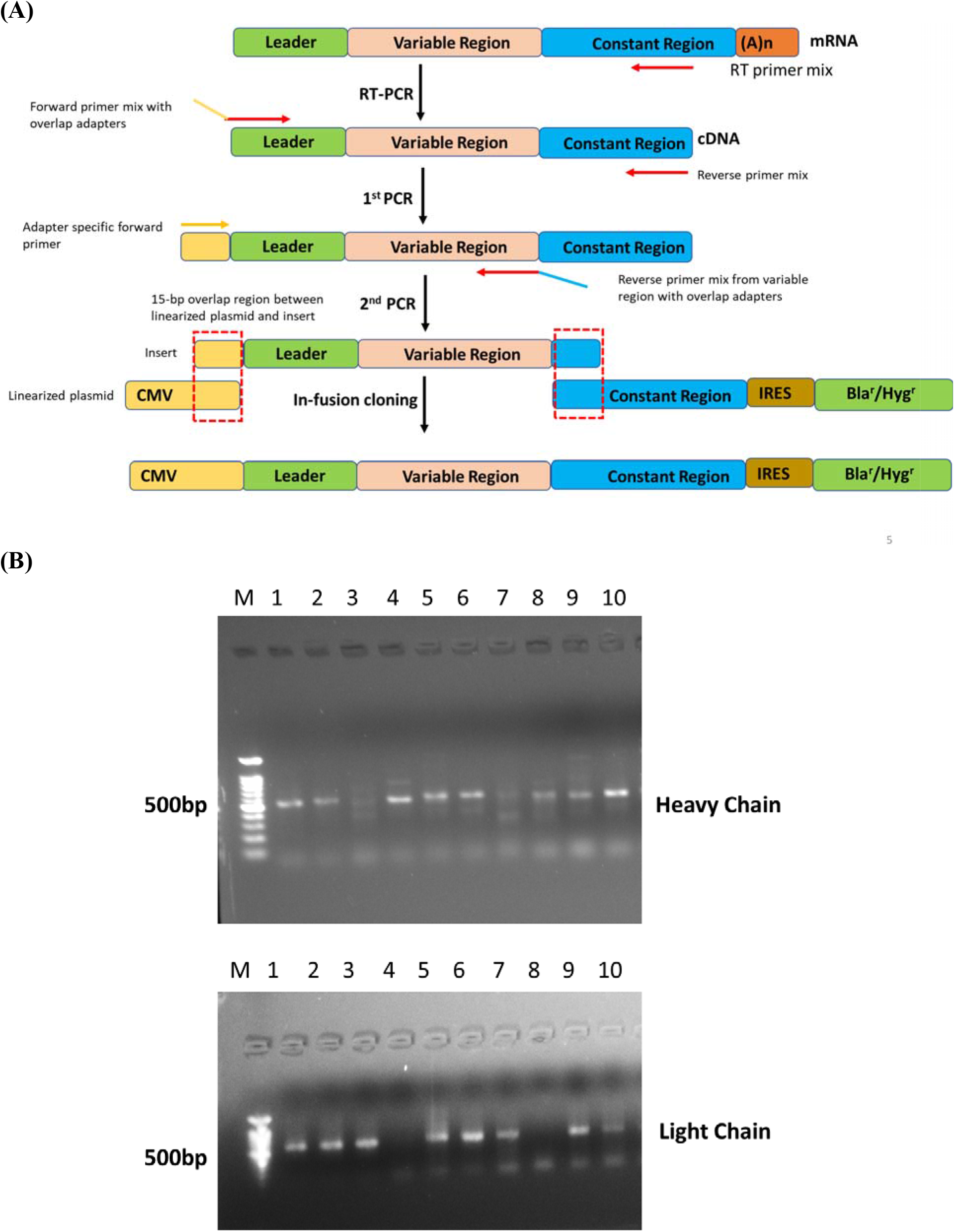
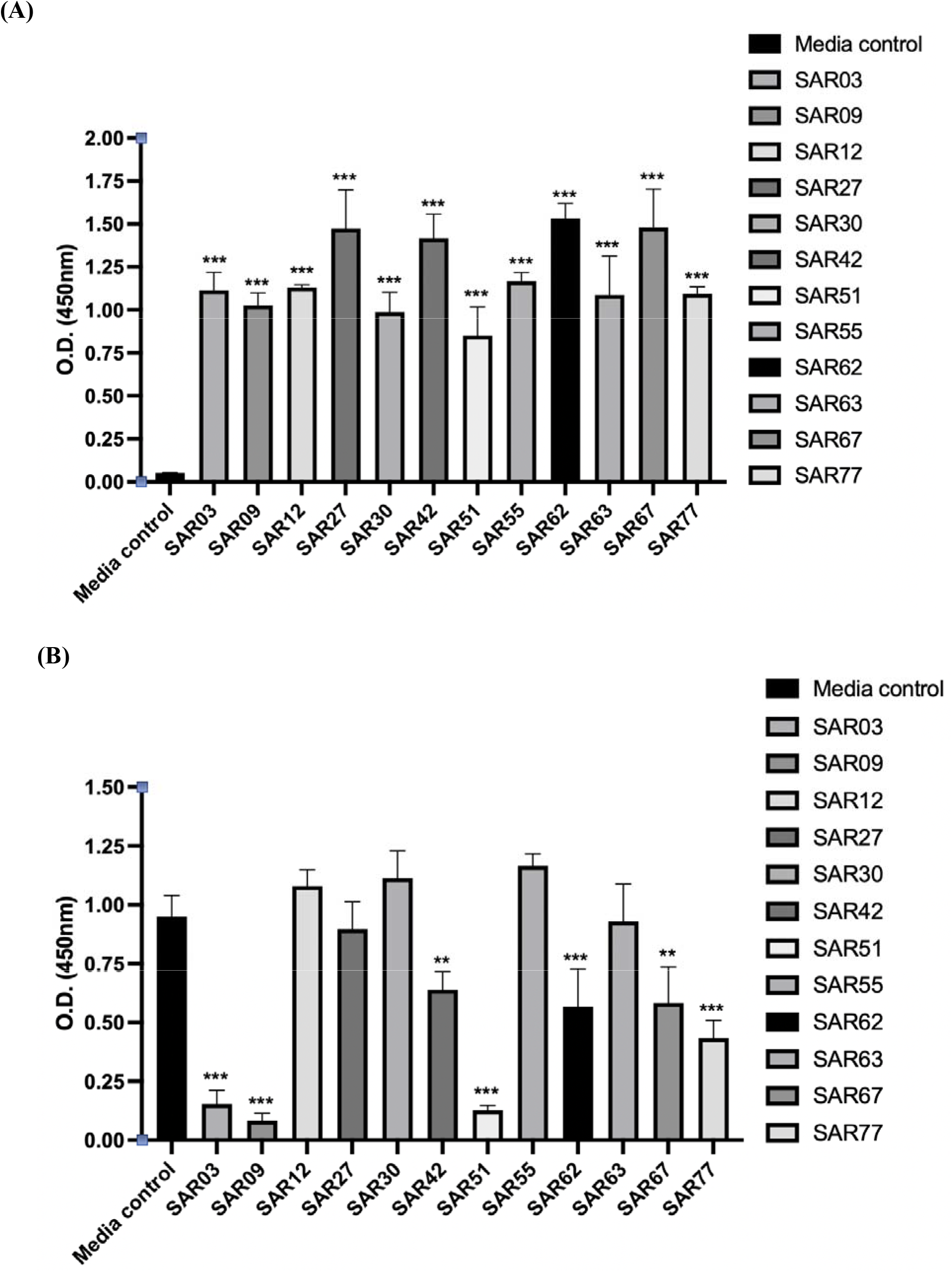
Binding analysis and neutralization activity of 12 antibody clones. **(A)** Cell culture supernatant from the cells transfected with heavy and light chain expressing plasmid was added to ELISA plates coated with recombinant RBD at 1/10 dilution. Anti-Human-HRP secondary antibody was used for ELISA detection. **(B)** Neutralization activity of cell culture supernatant was determined through competitive ELISA. Results are represented as Mean ± SD (n = 3). ***P<0.001, **P<0.01 on comparison cell culture supernatant with media control.

### 3.4 Generation of the stable cell line, antigen affinity, and neutralizing activity of potent antibodies

Freestyle-293F cells were transfected with a plasmid expressing heavy and light chain for three neutralizing clones. Transfected cells were then selected in presence of both hygromycin and blasticidin to generate stable clones expressing both chains. Limited dilution method was then used to generate monoclones and the clone showing the highest titer was selected for further characterization. High expressing stable Freestyle-293F monoclone was adapted to suspension culture and the supernatant was collected for recombinant antibody purification. Antibodies were purified through binding to Protein G agarose. Purified antibodies were first tested for their activity to neutralize Sars-CoV-2 pseudovirus binding to ACE-2 receptor. Recombinant antibodies at different concentrations were pre-incubated with Sars-CoV-2 pseudovirus and later added to HEK293 cells overexpressing the ACE-2 receptor. Percentage inhibition and IC50 values were calculated for top clones. SAR03 showed potent neutralizing activity (IC50-1.87 nM), while SAR09 (IC50-21.23 nM) and SAR051 (IC50-20.64 nM) showed slightly lower neutralizing activity **(Fig.5A).** SAR03 was also tested for its affinity to recombinant RBD protein through surface plasmon resonance (SPR). SAR03 showed potent affinity to RBD as evident from Kd-1.2 nM **(Fig.5B).** In our studied experiments, we finally identified SAR03 as the best mAb compared to other monoclones in terms of high-affinity capability and potent neutralization ability against SARS-CoV-2.

**Fig.5.**
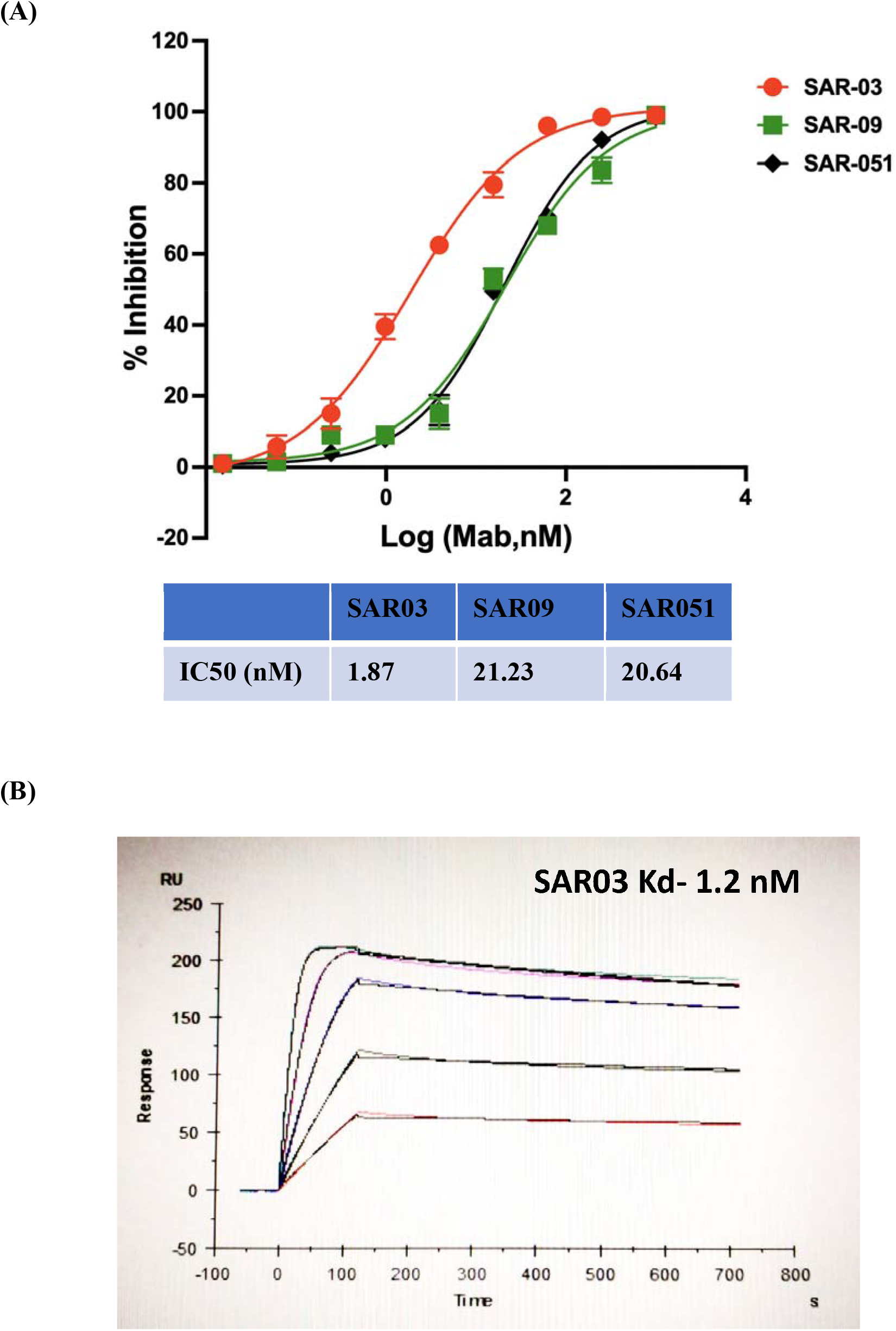
Determination of neutralization activity of top clones in pseudovirus reporter assay and affinity through SPR. **(A)** ACE-2 overexpressing cells were treated with Sars-CoV-2 pseudovirus in presence of different concentrations of purified antibodies. Luciferase activity was used as a measure of neutralizing activity. **(B)** SAR03 affinity against RBD was determined through SPR displayed on sensorgram plotted binding response (Response Unit; Ru) vs time (s).

## 4. Discussion

We describe here the isolation and sequencing of a high affinity and neutralizing mab against Sars-CoV-2 virus derived from a convalescent COVID-19 patient’s blood through an optimized, high-throughput, and efficient single B-cell cloning.

COVID-19 pandemic has impacted the world through its devastating affected human health either directly or indirectly. This pandemic has brought the foremost attention that focuses on the development of vaccines, novel antiviral agents, and mAbs. Furthermore, the establishment of neutralizing mAbs to SARS-CoV-2 exhibits great potential for both therapeutic and prophylactic applications and can offer a framework for vaccine design and development [25]. Numerous research groups have isolated mAbs through sorting of RBD antigen-specific B cells of infected patients who have recently recovered from SARS-CoV-2 using a single B-cell cloning [12, 13, 16–19]. The main target of SARS-CoV-2 neutralizing mAbs is the RBD domain of S1 surface glycoprotein (S1 protein) that mediates viral entry into host cells through interaction with ACE-2 receptors found on numerous cell types. Most studies conducted on mAbs isolated to date specifically target the RBD domain present on the spike protein. In general, for isolation of neutralizing antibodies through a single B-cell cloning method, blood samples were collected from patients recovered from SASR-CoV-2 infection. Furthermore, the RBD antigen-specific B cells are isolated from PBMC through staining with specific fluorescently labeled antibodies and sorting through FACS. Paired heavy and light chain sequences of Ab genes are then attained either using PCR or directly through a high-throughput single-cell sequencing [16, 26, 27]. In our study, we obtained the paired heavy and light chain genes using a rapid and high-throughput single B-cell screening platform for neutralizing Abs isolation and evaluated on the RT PCR platform that corroborated with the previous report [16].

We started with sorting of Sars-CoV-2 specific memory B-cells from convalescent patient’s blood using recombinant RBD protein as bait. Based on existing evidence that RBD-specific neutralizing antibodies can inhibit viral entry [22], the RBD domain was selected as the target antigen for memory B-cell sorting for the isolation of heavy and light chains. Using a smart gating strategy, we were able to sort memory B-cells specific for Sars-CoV-2 antigen. We found promising results for both binding and competitive (neutralization) ELISA conducted on patients’ plasma exhibit its high affinity towards RBD protein for the former and neutralize binding to recombinant ACE-2 protein for the latter Both of these initial studies suggested that the neutralizing activity of antibodies present in plasma can be attributed to the RBD domain, which might basically because RBD is composed with ACE-2 binding epitopes [28].

After sorting the cells in a single cell format, we used our in-house optimized RT-PCR and smart-PCR platform for rapid amplification of the paired variable region of the heavy and light chain of antibodies. cDNA was prepared in the same wells in a 96-well plate where the single cells were sorted. For heavy and light chains, RT-PCR primers (HC-RT and LC-RT) were used from a conserved region in the constant region. After RT-PCR, cDNA was divided into two plates for amplification of heavy and light chains separately. For amplification and cloning of variable regions, a two-step PCR using modified primer pairs was used. For the first PCR reaction, a mix of forward primer was designed from the 5’ region of the native signal peptide of the antibody clones. A unique adaptor sequence was also added at 5’ of the forward primer to be used as a forward primer in the second PCR step and for in-fusion cloning into expression plasmids. In the second PCR step, adaptor sequence was used as the forward primer and a mix of reverse primers, also containing a unique adaptor sequence, from the J-region were used to specifically amplify variable regions. B-cells showing a band for both heavy and light chain sequences were taken forward for cloning **(Fig.3B).** The adaptor sequences in the forward and reverse primers were homologous to the overhangs of linearized expression plasmids. This overlap was utilized to clone the antibody genes into the expression plasmids through in-fusion cloning. The ligation mix was transformed to DH5-α competent cells in 2-ml deep well plates (total 1ml of transformed mix). 500μl of the transformed mix was cultured overnight at 37°C and the plasmid was purified the next day. The remaining 500μl transformed mix was plated in LB-ampicillin plates for generating colonies. The plasmid was purified and transfected to freestyle-293F cells in a 96-well plate. 48-hours later, cell culture supernatant was tested for binding and neutralization activity through ELISA. For the positive clones showing binding and neutralization, DH5-α colonies on LB agar with ampicillin (100μg/ml) plates were then screened for a single clone (through colony PCR) containing the heavy and light chain sequences for large-scale plasmid purification and sequencing.

A stable cell line was generated for the final three clones in Freestyle-293F cells by transfecting the heavy and light chain expressing plasmids and selection using hygromycin and blasticidin. Recombinant antibodies were then purified from the supernatant of stable cells adapted to suspension culture. Purified proteins were then tested for neutralization activity through Sars-CoV-2 pseudovirus-based reporter assay. This pseudovirus is added to HEK293 cells, overexpressing the ACE-2 receptor, in presence of different concentrations of antibodies, and incubated for 72 hours. After the completion of incubation time, luciferase substrate was added, and relative luminescence activity was captured as an indicator of neutralizing activity. We could observe potent neutralizing activity with one of the clones, SAR03, while low activity was observed for the remaining two clones. The affinity of SAR03 towards was also assessed with SPR and potent affinity (Kd-1.2 nM) was observed for recombinant RBD protein.

In conclusion, we have successfully established an efficient and rapid method of screening neutralizing antibodies directly from recovered patients’ blood through single memory B-cell cloning. This method can be employed for isolating neutralizing antibodies against emerging infectious diseases in the future.

## Abbreviations

Sars-CoV-2: Severe Acute Respiratory Syndrome Coronavirus 2
RBD: Receptor-binding domain
ELISA: Enzyme-linked immunosorbent assay
SPR: Surface plasmon resonance
mAbs: Monoclonal antibodies
BCR: B cell receptor
PBMC: Peripheral blood mononuclear cells
ACE-2: Angiotensin-converting enzyme 2
IgG: Immunoglobulin
FACS: Fluorescence-activated cell sorting
PCR: Polymerase chain reaction

## Ethics Statement

Patient blood and PBMCs were provided by VB Pharma private limited, Gulbarga. The study was approved by the ethics committee of VB pharma private limited. Certified consents were obtained from all participants.

## Acknowledgments

Vinod Kumar Bagdu (VB Pharma) for providing patient blood and supporting the isolation of PBMCs; Sumanth VP, Anagha SM, Nidhi Singh-Supported the reagent preparation, assay development, and scientific inputs

## Authors Contributions

**PP, PB:** Concept, designed experimental work plan, performed experiments, cloning, stable cell line generation, cell culture, antibody binding assay, Data interpretation, conclusion, Manuscript writing; **PB, SS, NA, PA:** Performed experiments, B-cell sorting by FACS and analysis, antibody production, and purification, Neutralization assay, ELISA, Data acquisition, collection, analysis; **AS, MK**: Technical support, Data analysis, manuscript writing, final revision of the manuscript

